# Impact of white spotting alleles, including *W20*, on phenotype in the American Paint Horse

**DOI:** 10.1101/678052

**Authors:** Samantha A. Brooks, Katelyn M. Palermo, Alisha Kahn, Jessica Hein

## Abstract

The American Paint Horse Association (APHA) records pedigree and performance information for their breed, a stock-type horse valued as a working farm or ranch horse and as pleasure horses. As the name implies, the breed is also valued for attractive white spotting patterns on the coat. The APHA utilizes visual inspections of photographs to determine if coat spotting exceeds threshold anatomical landmarks considered characteristic of desirable patterns. Horses with sufficient white patterning enter the “Regular” registry, rather than the “Solid Paint-bred” division, providing a threshold modeled phenotype. Genetic studies previously defined sequence variants corresponding to 35 alleles for white spotting in the horse. Here, we calculate the allele frequency for nine common white spotting alleles in the American Paint horse using a sample of 1,054 registered animals. The APHA spotting phenotype is altered by additive interactions among spotting loci, and epistatically by the *MC1R* and *ASIP* genes controlling pigment production. The *W20* allele within the *KIT* gene, independent of other known spotting alleles, was strongly associated with the APHA-defined white spotting phenotype (p = 1.86 x10^−18^), refuting reports that *W20* acts only as a modifier of other underlying white spotting patterns. The parentage of an individual horse, either American Paint or American Quarter Horse, did not alter the likelihood of entering the APHA Regular registry. An empirical definition of the action of these genetic loci on the APHA-defined white spotting phenotype will allow more accurate application of genome-assisted selection for improving color production and marketability of APHA horses.

## Introduction

Founded in 1962, the American Paint Horse Association’s mission includes preserving the lineage of colorful, versatile stock horses through registration with the association (Hood 1987). Since its inception, registration of the Paint Horse has been based primarily on a combination of pedigree and phenotype in order to select for the white-patterned, stock-type horse valued by APHA’s founders—this includes acceptance of “cropout” Quarter Horses, who were refused registration with the American Quarter Horse Association until 2004 (AQHA 2019). As of January 1, 1980, the association requires all fully-registered APHA horses to have a sire and dam registered with either the American Paint Horse Association, the American Quarter Horse Association or The Jockey Club, a standard that remains today (APHA 2018). With more than 1.09 million horses registered with its organization as of May 2019, the APHA ranks as the second-largest equine breed association in the world. In the US alone, the equine industry contributes a total of approximately $122 billion to the U.S. economy and 1.7 million jobs (American Horse Council Foundation 2019). While the American Paint Horse is valued for conformation and performance traits, the white spotting pattern on the coat exerts an overwhelming influence on the economic value of individual horses within this large breed registry (Brooks et al. 2007).

During the APHA registration process, horses are designated into one of two sub-registries based on the presence or absence of qualifying natural white markings on their coats: the Regular registry or the Solid Paint-bred registry. This is done through phenotypic observation of photographs submitted of the horse, analyzed by trained APHA staff members against a set of thresholds (reference lines along the hocks, knees and head) determined by the association’s board of member-elected directors, not empirical scientific data. Regular registry status is granted for horses that meet white spotting requirements as outlined in the *APHA Rule Book* which has contained a diagram describing these thresholds since 1973 (APHA 2018). The patterns desired by the APHA extend beyond what is typically observed as an average white marking on the face or legs (Haase et al. 2013) and exclude markings created by the *Leopard* locus (Bellone et al. 2013).

In brief, unpigmented areas characterizing APHA-accepted white spotting patterns must include both white hair and skin extending beyond reference lines determined by the association’s elected governing body; those reference lines are not based on empirical thresholds of known white spotting patterns’ distribution zones. On the face, qualifying depigmented areas must extend beyond a reference line from the base of the horse’s ear to the outside corner of the eye to the corner of the mouth and under the chin to the opposite corner of the mouth (Figure 1). On the body, qualifying depigmented areas must occur above a perpendicular line around the leg located at the center of the knee or hock (Figure 1). These spotting patterns must include a minimum of two inches of solid white hair with some underlying unpigmented skin in the qualifying area. The Solid Paint-Bred registry is used for horses who meet the parentage requirement but lack the minimum requirement for white spotted skin and coats as defined in the rulebook (APHA 2018).

**Figure 1:**
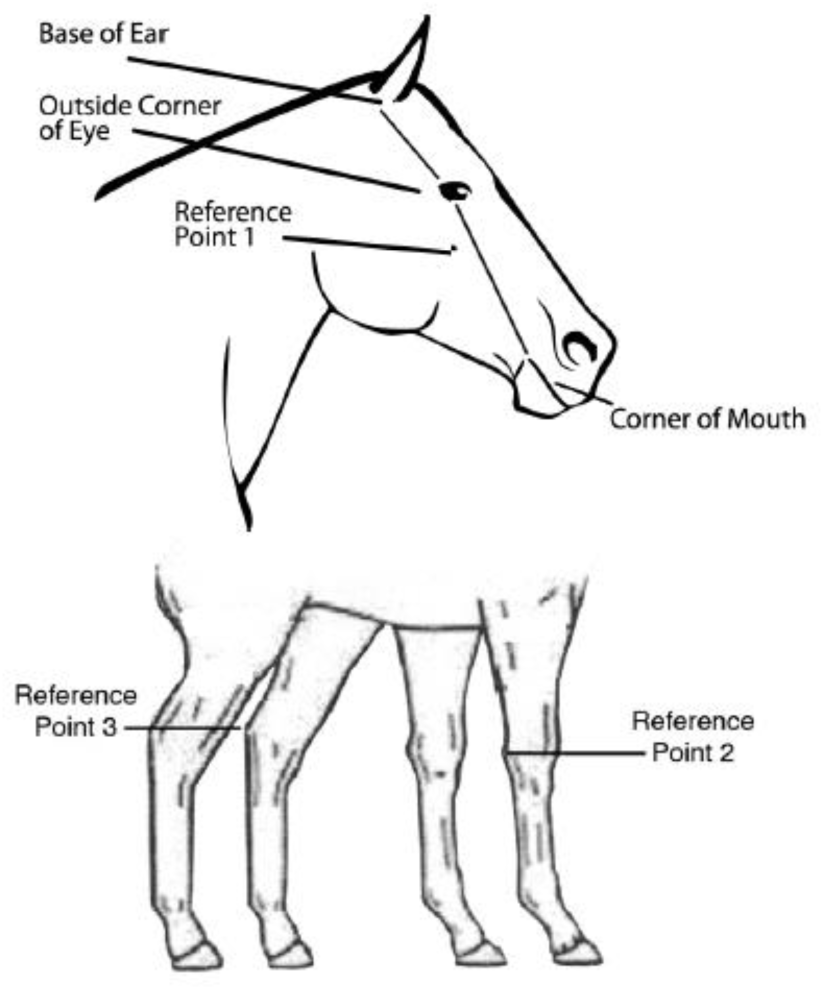
Anatomical landmarks on the face and limbs of the APHA horse are used to determine the extent of white spotting patterns, and to designate each registered horse to either the Regular or Solid Paint-bred sub-registries. To enter the Regular registry, a proposed horse must possess at least 2” of white hair, as well as underlying unpigmented skin, on the body surface beyond these reference lines.

**Figure 2:**
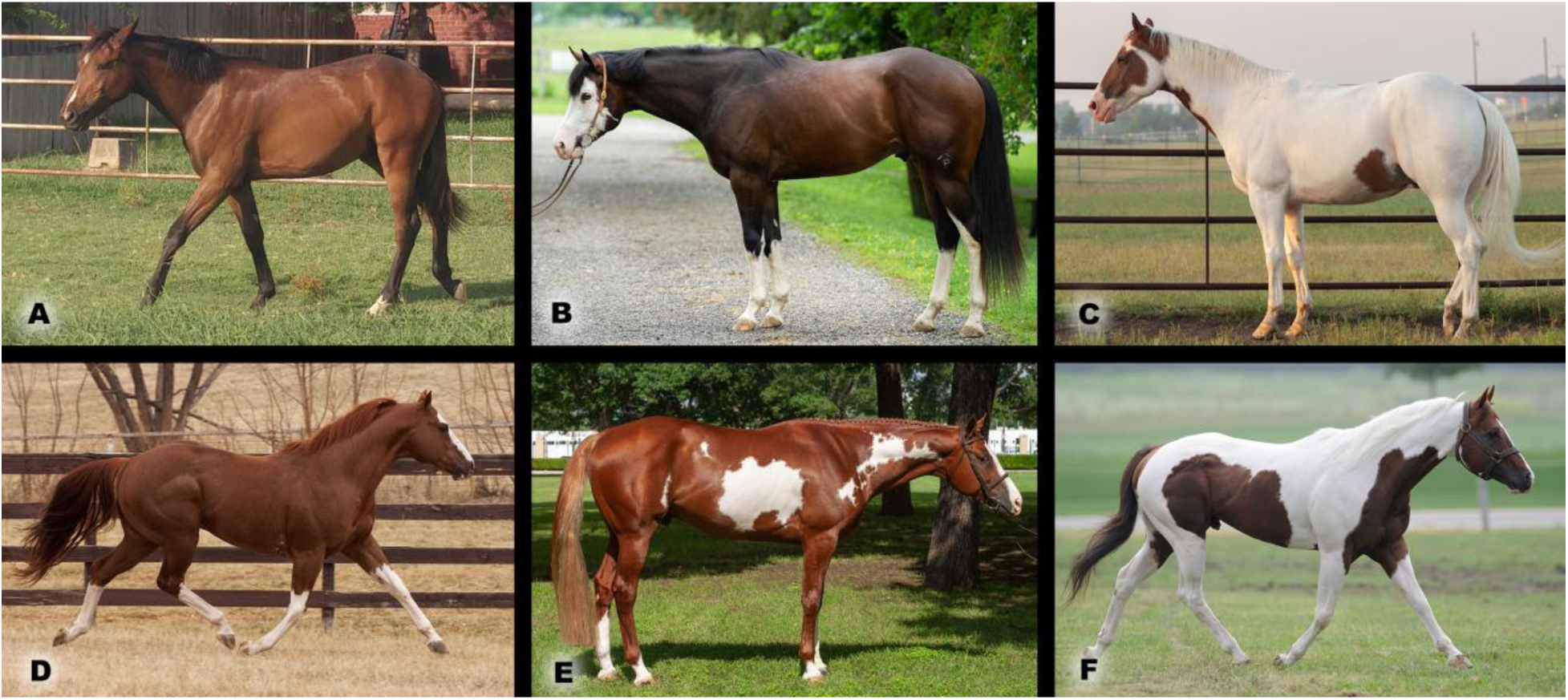
Diverse white spotting phenotypes are captured by the American Paint Horse Registry. Horse A is categorized in the Solid registry though he possesses the same *EDNRB*O* white spotting genotype as horse E. Horses B-F are all classified as Regular registry by photographic assessment (photo credits: J. Hein)

When registering a Paint Horse today, the applicant provides identifying information pertaining to the foal, including its date of birth, color, pattern, sex, parentage and ownership. The APHA requires parentage verification via genetic testing when the horse is the product of breeding with transported or frozen semen, embryo transfer or vitrified embryo transfer, or other special situations, such as when registering a horse over age 10, and for all Quarter Horses and Thoroughbreds applying for registration as Paint Horses (APHA 2018).

The applicant also provides a minimum of four full-color photographs of the horse, showing the entire animal from the left side, right side, front and rear. Additional photographs of spotted areas of the coat might be required by APHA staff to verify that the pattern meets registration guidelines in terms of the location, size and requirement for underlying skin to also be unpigmented (APHA 2018). The photographs become part of the horse’s permanent record at APHA and are used to identify the animal throughout its life. Trained APHA staff members determine the registration category (Solid Paint-Bred Registry or Regular Registry for each horse by visual inspection to confirm that a horse’s natural white spotting patterns meets APHA’s standard. The quality of the submitted photographs is variable, adding additional challenges to the interpretation of photographic evidence of the pattern relative to the APHA standards.

In the horse, at least 35 known white spotting polymorphisms underly white spotting patterns on the skin and coat: for a comprehensive and current listing of spotting alleles in the horse see OMIA.org (OMIA 2019). Of these known alleles, nine are common to and routinely tested for in the APHA population (Table 1). Others, like *MITF-Splash White 4* and many of the *KIT*W* alleles, are breed specific and not present in the Paint Horse, while some like *MITF-Splash White 5* were not yet commercially available for testing at the time the study began and therefore were not examined (Henkel et al. 2019).

**Table 1:** Details of the nine polymorphisms genotyped for this analysis. The “Reference” allele refers to the EquCab3.0 assembly and also happens to be the presumed ancestral or wild type for each of these nine variants.

| Locus | Reference Allele | Alternate Allele | Phenotype | Inheritance Pattern for Alternate Allele | Reference |
| --- | --- | --- | --- | --- | --- |
| *EDNRB* | *N* * | *O* | Overo spotting/OLWS | Dominant, homozygous lethal | Metallinos *et al.* 1998; Santschi *et al.* 1998; Yang *et al.* 1998 |
| *ECA3* | *N* * | *TO* | Tobiano spotting | Dominant | Brooks *et al.* 2007 |
| *KIT* | *N* * | *SB1* | Sabino spotting | Additive | Brooks & Bailey 2005 |
| *KIT* | *N* * | *W5* | Variable white spotting | Dominant, presumed homozygous lethal | Haase *et al.* 2009 |
| *KIT* | *N* * | *W10* | Variable white spotting | Dominant, presumed homozygous lethal | Haase *et al.* 2009 |
| *KIT* | *N* * | *W20* | Variable white spotting | Dominant | Hauswirth *et al.* 2013 |
| *MITF* | *N* * | *SW1* | Splashed white spotting | Dominant | Hauswirth *et al.* 2012 |
| *MITF* | *N* * | *SW3* | Splashed white spotting | Dominant, presumed homozygous lethal | Hauswirth *et al.* 2012 |
| *PAX3* | *N* * | *SW2* | Splashed white spotting | Dominant | Hauswirth *et al.* 2012 |
| *ASIP* | *A* | *a* | Black/Bay pigment distribution | Recessive | Rieder *et al.* 2001 |
| *MC1R* | *E* | *e* | Black/Red pigmentation | Recessive | Marklund *et al.* 1996 |

\*For these loci the “*N*” designation is used by the UC Davis commercial testing service to communicate a wild type allele.

Among the APHA community, significant controversy surrounds the action of *KIT*W20*. The first published observation of the *KIT*W20* allele came from a study searching for causative variants in horse with white coats, phenotypes that left the majority of the skin surface totally unpigmented, and therefore did not attempt to document relatively less extensive white spotting phenotypes among their sampled population (Haase et al. 2007). Subsequently, the authors examined the *KIT*W20* variant in a larger set of horses (n= 52) and noted a compound heterozygote effect creating extensive white patterning in horses with the *KIT*W5/W20* genotype, and observing a significant effect of the *KIT*W20* allele generating a white spotting phenotype more expansive than the typical white markings on the face and legs (Hauswirth et al. 2013). These authors state the circumstances in their publication: “We previously discovered a missense variant in exon 14 of the equine *KIT* gene (c.2045G>A; p.Arg682His) but did not immediately recognise [sic] its functional role (Haase et al. 2007).” However, in the eyes of the lay audience, the latter publication was eclipsed by the 2007 paper containing limited observations, and the lack of data specifically addressing the phenotype produced by the *KIT*W20* allele led to public claims that there is no, or only a “subtle” impact from the *KIT*W20* allele on white spotting (https://www.vgl.ucdavis.edu/services/horse/dominantwhite.php).

As a whole, the population frequencies of all white-spotting alleles lack investigation, and despite the easily applied Mendelian inheritance patterns for most of these alleles, genetic testing for selection of breeding stock remains underutilized in the industry. Only recently has the APHA industry expressed an interest in using readily available genetic tools to improve their color production. Previously, selective breeding for spotting patterns in the American Paint Horse was commonly based on hearsay, personal experience, and limited observations. Starting in 2017, the APHA has developed rules that help designate a horse to the Regular Registry based on presence of white-spotting alleles. These new rules give breeders and horse owners additional tools to make responsible breeding decisions in an effort to produce the healthiest, most marketable horse possible. This more inclusive approach encourages objective registration of foals with phenotypes not well characterized by the APHA threshold method alone, adding significant economic value to these horses, which in turn can ultimately improve their long-term welfare (Hein 2017).Recent work highlighting “exceptions” to proposed qualitative phenotypes for these loci emphasizes the need for application of impartial and quantitative assessments capturing the full breadth of phenotypes resulting from these variants (Druml et al. 2018). Toward this ideal, the goals of this study were to: establish allele frequencies for known white spotting alleles in the American Paint Horse, investigate association of these loci with a threshold phenotype uniformly defined by the APHA, identify interactions between loci, any effect due to the sex of the horse and to test association of the *W20* allele with the APHA photo-based threshold phenotype.

## Materials and Methods

### Retrospective Registration Records

The American Paint Horse Association provided data for a sampling of 1,054 horses born and registered between 1992 and 2018 including: registered name and number, registration type (“Regular” registry *i.e.* possessing a spotted coat or “Solid Paint-Bred” phenotype), age, and sex for each horse, as well as the registry and registration type of the sire and dam for that horse. The original photos submitted by the applicant for evaluation of the white spotting pattern on the horse at the time of registration were also provided by the APHA and used to confirm predicted phenotypes by visual inspection from a single experienced observer blinded to the identity and genotype of the horse (SAB). In 2012, the APHA began partnering with commercial providers of genetic testing to record genotypes on their registered horses. These 1,054 horses were voluntarily tested for color alleles by their owners for their own educational use or interests. Results were already on file with APHA and although they are not a randomly extracted subset of all APHA registrations, the large sample size does offer the opportunity to investigate important questions regarding the action of these alleles.

### Genotypes

Each horse was genotyped for nine spotting pattern loci using a commercial service (Veterinary Genetics Laboratory, University of California, Davis) and the data shared with the APHA through a collaborative agreement. Alleles at the following loci were examined (Table 1): *KIT-TO, SB1, W5, W10, W20* or “*N”* (Brooks & Bailey 2005; Brooks *et al.* 2007; Haase et al. 2009; Hauswirth *et al.* 2013), *EDNRB-O* or “*N”* (Metallinos et al. 1998; Santschi et al. 1998; Yang et al. 1998)*, MITF-SW1, SW3* or *N* and *PAX3-SW2* or “*N”* (Hauswirth et al. 2012). The testing service provider designates the wild type allele as “*N”.* Additional genetic variants associated with white spotting patterns in the horse are documented in the scientific literature, but were either not available commercially at the time of testing and/or do not commonly segregate in the APHA and are therefore not relevant to this population (for a comprehensive catalog of spotting alleles in the horse see OMIA.org). Additionally, two known pigmentation loci were genotyped for investigation of epistasis: *MC1R-E* or *e* (Marklund et al. 1996)*, ASIP-A* or *a* (Rieder et al. 2001). For the purpose of calculating allele frequencies we refer to the allele carried by the reference genome assembly (EquCab3.0) as the wild type (Kalbfleisch et al. 2018).

### Statistical analyses

Analyses were conducted in the JMP Pro v14.1.0 package (SAS Institute Inc.). The additive effect of multiple white spotting loci was tested by a logistic regression of the number of alleles against the two possible registry categories. Linkage, assessed as LOD scores between coat color loci on ECA3, was assessed using HaploView v4.2 (Barrett 2009). To measure the impact of the *MC1R-ASIP* signaling system on white spotting, a subset of 364 horses that possessed only one spotting allele (controlling for any effect due to interactions between multiple spotting loci) was utilized, excluding all horses possessing a *Tobiano* allele (in order to avoid the impact of linkage between *MC1R* alleles and *TO).* Given the well documented interaction between alleles at *ASIP* and its antagonistic target, the *MC1R* receptor, a logistic regression model was constructed comparing the APHA registry phenotype (Regular or Solid Paint-Bred) with the linear ranking of the genotypes by the predicted signaling activity and base color phenotype. Thus, the *E/-a/a* genotype, likely resulting in a constitutively active *MC1R* receptor and the black base color, was scored “0”; the wild-type genotype *E/-A/-*, predicted to have normal signaling activity and a bay base color, was given a “1”; and the *e/e* genotype, resulting in a loss of *MC1R* signaling regardless of the *ASIP* genotype and a chestnut base coat, received a score of “2” (Barrett et al. 2005). Association between genotypes for the *W20* allele and the APHA threshold phenotype was assessed in a subset 529 horses bearing no other white spotting alleles at any of the other eight sites tested and using a Chi-square test under a dominant model (merging the *W20/N* and *W20/W20* genotypes).

## Results and Discussion

### Phenotype distribution varies for each white spotting genotype

The 1,054 horses registered with the APHA in the examined time-period included 406 stallions, 90 geldings and 558 mares submitted for registration at a mean age of 1.07 years between 1992 and 2018. 90% of the sample comprised horses registered post-2004. Among these 1,054 horses, 773 were designated to the Regular registry, while 281 entered the Solid Paint-Bred category. This sampling did not represent an unbiased observation of all foals produced from APHA-registered breedings. Not all foals are submitted for registration (especially if the foal is unlikely to achieve Regular registry status), and because DNA genetic testing for white spotting alleles is not required for registration, the available data for this retrospective study is limited to horses registered with APHA that also had DNA testing results available for investigation. Records are not retained by the APHA on submitted horses ultimately rejected for registration. Nonetheless, this sample is the largest collection of genotype data and white spotting phenotypes in the horse.

Genotypic and allele frequencies for each of the nine variants examined are presented in Table 2. Presence of just a single alternate allele resulted in APHA designation to the Regular registry for *KIT*TO*, *MITF*SW3*, and *KIT*SB1*, although the latter two genotypes were only rarely observed (n = 3 and 8 respectively, Table 3). As expected, we did not observe a homozygous *EDNRB*O* genotype as this causes the well-documented Lethal White Overo Syndrome (Metallinos *et al.* 1998; Santschi *et al.* 1998; Yang *et al.* 1998). However, a single horse homozygous for the *SW2* allele was identified, which is a state previously hypothesized to be lethal based on comparisons to similar variants in the *PAX3* gene of other species (Hauswirth *et al.* 2012). The *SW2/SW2* genotype was confirmed in a second DNA sample submitted independently and anonymously. Veterinary examinations of this horse were unavailable at the time of writing, but his owner reports that aside from apparent deafness she is healthy and strong (Figure 3.).

**Table 2.** Genotype counts and allele frequencies for nine white spotting variants and two base coat color loci genotyped among 1054 APHA registered horses, within each of the two registration categories. Data are presented as genotype counts for the homozygous reference (Hom Ref), heterozygous (Het) and homozygous alternate (Hom Alt) states, as well as the frequency of the reference (*f*Ref) and alternate (*f*Alt) alleles.

|  | Regular Registry<br>(n= 773) |  |  |  |  | Solid Paint-bred Registry<br>(n= 281) |  |  |  |  | Total<br>(n= 1054) |  |  |  |  |
| --- | --- | --- | --- | --- | --- | --- | --- | --- | --- | --- | --- | --- | --- | --- | --- |
|  | Hom<br>Ref | Het | Hom<br>Alt | *f*Ref | *f*Alt | Hom<br>Ref | Het | Hom<br>Alt | *f*Ref | *f*Alt | Hom<br>Ref | Het | Hom<br>Alt | *f*Ref | *f*Alt |
| *EDNRB* * *O*<br>(Frame Overo) | 570 | 203 | 0 | 0.87 | 0.13 | 263 | 18 | 0 | 0.97 | 0.06 | 833 | 221 | 0 | 0.88 | 0.12 |
| *TO*<br>(Tobiano) | 572 | 142 | 59 | 0.83 | 0.17 | 281 | 0 | 0 | 1.00 | 0.00 | 853 | 142 | 59 | 0.88 | 0.12 |
| *KIT* * *SB1*<br>(Sabino1) | 748 | 24 | 1 | 0.98 | 0.02 | 281 | 0 | 0 | 1.00 | 0.00 | 1029 | 24 | 1 | 0.99 | 0.01 |
| *KIT* * *W05*<br>(Variable White) | 773 | 0 | 0 | 1.00 | 0.00 | 281 | 0 | 0 | 1.00 | 0.00 | 1054 | 0 | 0 | 1.00 | 0.00 |
| *KIT* * *W10*<br>(Variable White) | 771 | 2 | 0 | 1.00 | 0.00 | 281 | 0 | 0 | 1.00 | 0.00 | 1052 | 2 | 0 | 1.00 | 0.00 |
| *KIT* * *W20*<br>(Variable White) | 404 | 319 | 50 | 0.73 | 0.27 | 196 | 74 | 11 | 0.83 | 0.17 | 600 | 393 | 61 | 0.76 | 0.24 |
| *PAX3* * *SW2*<br>(Splashed<br>White) | 757 | 15 | 1 | 0.99 | 0.01 | 272 | 9 | 0 | 0.98 | 0.03 | 1029 | 24 | 1 | 0.99 | 0.01 |
| *MITF* * *SW1*<br>(Splashed<br>White) | 680 | 90 | 3 | 0.94 | 0.06 | 263 | 17 | 1 | 0.96 | 0.07 | 943 | 107 | 4 | 0.95 | 0.05 |
| *MITF* * *SW3*<br>(Splashed<br>White) | 769 | 4 | 0 | 0.99 | 0.01 | 281 | 0 | 0 | 1.00 | 0.00 | 1050 | 4 | 0 | 1.00 | 0.00 |
| *ASIP* * *A*<br>(Bay/Black) | 316 | 323 | 134 | 0.62 | 0.38 | 115 | 118 | 48 | 0.62 | 0.38 | 431 | 441 | 182 | 0.62 | 0.38 |
| *MC1R* * *E*<br>(Black/Red) | 78 | 234 | 461 | 0.25 | 0.75 | 9 | 82 | 190 | 0.16 | 0.82 | 87 | 316 | 651 | 0.23 | 0.77 |

**Table 3:**
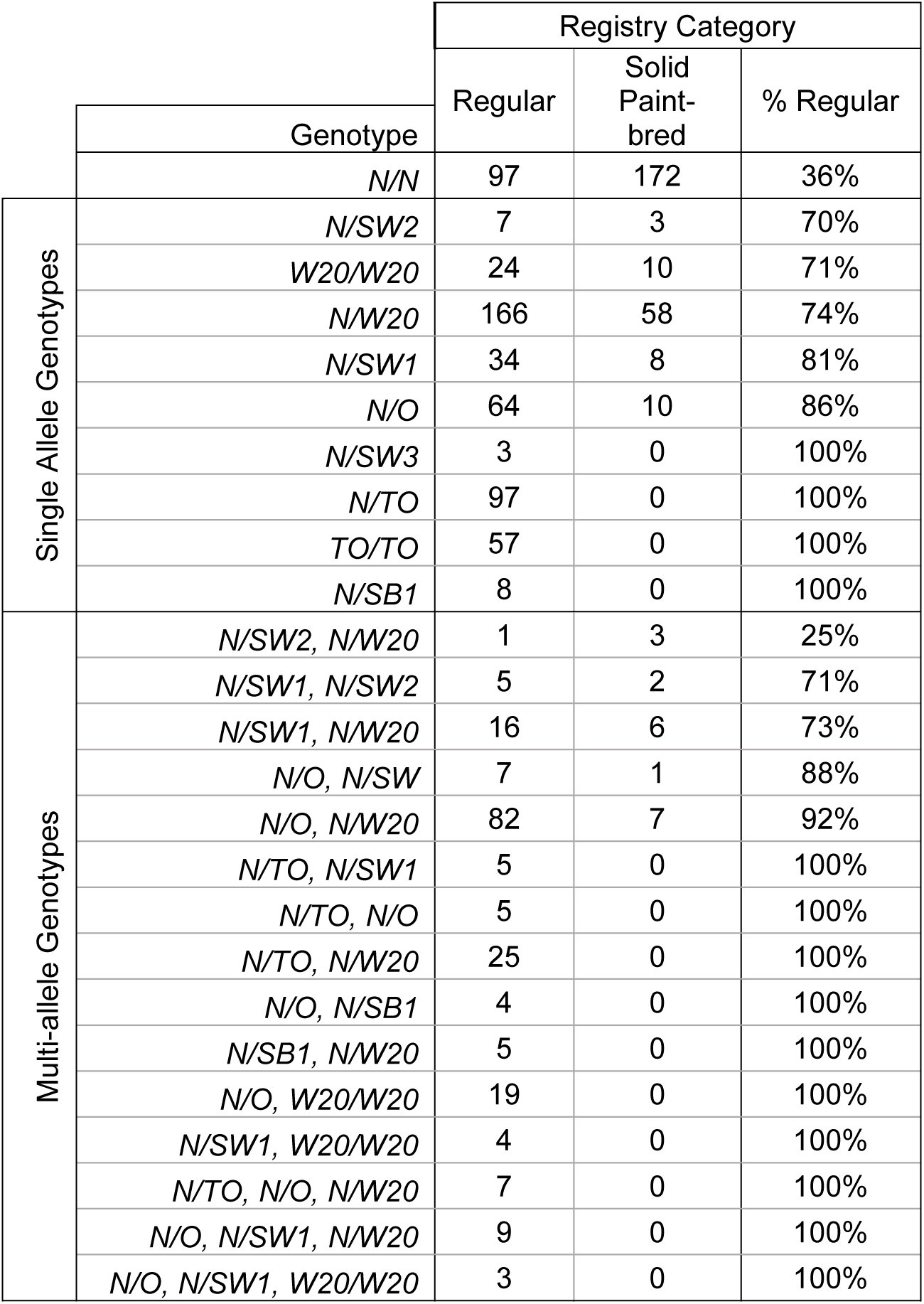
Horse designation to the Regular or Solid Paint-bred registry divisions by genotype and genotype combinations (for simplicity, the table reports only genotypes/combinations observed in at least three horses.)

|  |  | Registry Category |  |  |
| --- | --- | --- | --- | --- |
|  |  | Regular | Solid Paint-bred | % Regular |
|  | Genotype |  |  |  |
|  | *N/N* | 97 | 172 | 36% |
| Single Allele Genotypes | *N/SW2* | 7 | 3 | 70% |
|  | *W20/W20* | 24 | 10 | 71% |
|  | *N/W20* | 166 | 58 | 74% |
|  | *N/SW1* | 34 | 8 | 81% |
|  | *N/O* | 64 | 10 | 86% |
|  | *N/SW3* | 3 | 0 | 100% |
|  | *N/TO* | 97 | 0 | 100% |
|  | *TO/TO* | 57 | 0 | 100% |
|  | *N/SB1* | 8 | 0 | 100% |
| Multi-allele Genotypes | *N/SW2, N/W20* | 1 | 3 | 25% |
|  | *N/SW1, N/SW2* | 5 | 2 | 71% |
|  | *N/SW1, N/W20* | 16 | 6 | 73% |
|  | *N/O, N/SW* | 7 | 1 | 88% |
|  | *N/O, N/W20* | 82 | 7 | 92% |
|  | *N/TO, N/SW1* | 5 | 0 | 100% |
|  | *N/TO, N/O* | 5 | 0 | 100% |
|  | *N/TO, N/W20* | 25 | 0 | 100% |
|  | *N/O, N/SB1* | 4 | 0 | 100% |
|  | *N/SB1, N/W20* | 5 | 0 | 100% |
|  | *N/O, W20/W20* | 19 | 0 | 100% |
|  | *N/SW1, W20/W20* | 4 | 0 | 100% |
|  | *N/TO, N/O, N/W20* | 7 | 0 | 100% |
|  | *N/O, N/SW1, N/W20* | 9 | 0 | 100% |
|  | *N/O, N/SW1, W20/W20* | 3 | 0 | 100% |

**Figure 3:**
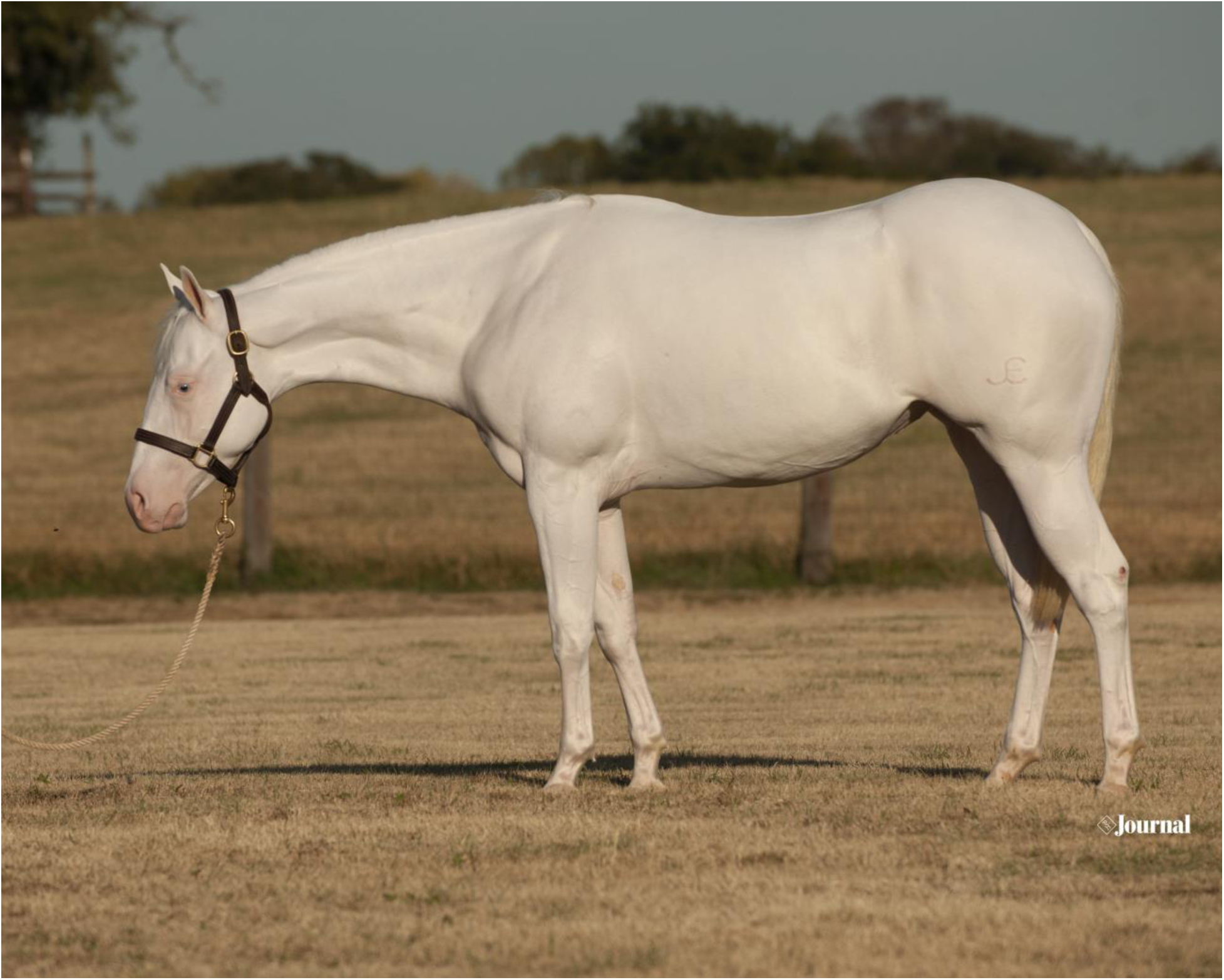
Previously thought to be lethal in the homozygous state, this carrier of the *PAX3 SW2/SW2* homozygous genotype has extensive depigmentation of the skin and coat (photo credit: Paint Horse Journal)

109 horses possessed at least one white spotting allele but were APHA registered Solid Paint-bred based on visual examination of photographs. Visual inspection of the photographs by a single blinded observer revealed that the extent and distribution of white markings in these horses often fell just below the guidelines of the *APHA Rule Book* (2018) but were more extensive than the common white markings described as “socks” on the limbs or a “blaze” on the face in previous studies (Haase *et al.* 2013). The phenotype of these horses likely represents the tail of the natural distribution of total depigmented area characteristic of the unique reduction in melanocyte migration for each of these alleles. Previous work observed production of foals with phenotypes very typical of the *TO* allele from parents with a *TO* allele but a very minimal white phenotype (Stachurska & Jansen 2015). Thus, these horses likely possess genetic value as producers of future spotted foals, although on the surface they do not fit the visual description of a Regular registry horse.

One horse categorized as Solid Paint-Bred possessed a total of five white spotting alleles and an entirely white coat. Although this animal has a high breeding value for white spotting, it was registered in the less-valued Solid Paint-Bred category because the horse did not exhibit at least two inches of contrasting colored hair in his coat, per APHA registration guidelines. In these cases, phenotyping by photograph and use of the *APHA Rule Book* description for Regular registry white spotting patterns misses horses with superior genetic value for production of white spotting patterns.

For 97 horses recorded by APHA in the Regular registry based on their photographic phenotype, genetic testing results did not identify possession of any of the nine white spotting alleles evaluated. In some cases, white spotting patterns in these horses may be due to rare genetic markers not tested in this study, or patterns for which genetic markers/causative variations are not yet known. Visual inspection of the registration photographs for these 97 horses revealed that 31 possessed patterns consistent with white spotting loci. These 31 included many phenotypes resembling those generated by alleles known but not genotyped in this cohort, including *MITF*SW4* (Hauswirth *et al.* 2013), *MITF*SW5* (Henkel *et al.* 2019), and more than 27 other alleles at the *KIT* locus. We also observed among these 31 horses some well-recognized patterns with yet unknown genetic etiology (*i.e.* the “Rabicano” pattern) and two horses with previously undocumented and distinct white spotting phenotypes. Phenotypes for the remaining 66 (68%) horses fell within the described range for common white markings of the face and legs (Haase *et al.* 2013), or typified the roan and grey coat colors. Roan and grey do introduce white hair into the coat, but in a more interspersed manner than desired by the APHA (Marklund et al. 1999; Sundstrom et al. 2012).

In total, the predicted phenotype from the genotypes at the nine spotting variants investigated here agreed with the Regular registry (spotted) classification in 95.4% of horses, demonstrating that these nine alleles account for an overwhelming majority of spotting phenotypes present in the APHA. However, the photo-based designation to the Regular or Solid Paint-Bred categories disagreed with the genotyped presence or absence of a white spotting allele in 17% of horses (206) submitted for registration. This discordance demonstrates the need for separate selection tools and interpretations for each of the functions of the registry: quantification of desirable visual phenotypes versus assessment of breeding value for production of white spotted foals.

### Cumulative interactions among spotting loci, and between spotting loci and alleles for base coat color, additively increase white spotting phenotypes above the APHA threshold

The frequency of the Regular registry threshold phenotype varied by allele and by allele combination (Table 3), however sample sizes within each possible combination were still too small to statistically examine specific interactions between loci. However, excluding the one horse possessing five alleles and an entirely white coat (considered a Solid Paint-Bred under current rules,) horses with an increasing number of spotting alleles are more likely to exhibit a phenotype that exceeds the APHA threshold for the Regular registry (Chi-square = 226.11, p < 0.0001, Table 4). Thus, the nine spotting alleles examined here act collectively in an additive fashion, increasing the proportion of white on the horse above the APHA Regular registry phenotype threshold.

**Table 4:** Horse counts for the Regular and Solid Paint-bred phenotypes compared to the total number of white spotting pattern alleles at the nine loci tested, and genotypes at the *MC1R* and *ASIP* coat color loci in a subset of 368 horses.

|  |  | Registry Category |  |  |
| --- | --- | --- | --- | --- |
|  |  | Regular | Solid Paint-bred | % Regular |
| Total # of White Spotting Alleles<br>(n= 1054) | 0 | 97 | 172 | 36% |
|  | 1 | 382 | 79 | 83% |
|  | 2 | 240 | 29 | 89% |
|  | 3 | 51 | 0 | 100% |
|  | 4 | 3 | 0 | 100% |
|  | 5 | 1 | 0 | 100% |
| *MC1R*<br>-*ASIP*<br>Geno.<br>(n=368) | *E*-, *a/a* | 13 | 7 | 65% |
|  | *E*-, *A*- | 57 | 23 | 71% |
|  | *e/e*, -/- | 219 | 49 | 82% |

Although the *MC1R* signaling pathway primarily contributes to pigment switching between eumelanin and pheomelanin, loss-of-function alleles at the *MC1R* locus can also reduce migration of melanocyte stem cells (Chou et al. 2013). Assessing the effect of MC1R signaling on depigmentation phenotypes resulting from the *KIT* locus is not straightforward as the physical proximity of these two genes on ECA3 could produce an association due to linkage. Indeed, suppression of recombination by the *Tobiano* allele, a 36 Mb paracentric inversion encompassing a 41 Mb span between the *KIT* and *MC1R* genes is well documented (Trommershausen-Smith 1978; Brooks *et al.* 2007). In this dataset, we observed strong linkage between *MC1R*E* and the *TO* inversion allele (n = 1054, LOD= 40.96), but not between either *MC1R* allele and *KIT*\**W20* (LOD= 0.05).

Before assessing modification of spotting phenotype by alleles at *MC1R* and *ASIP*, we selected a subset of horses that possessed just one spotting allele (avoiding any coincidental interaction across multiple spotting alleles) and excluding any horse with a *TO* allele (likely to produce association to *MC1R*E* due to recombination suppression generated by the *TO* inversion). Among these 368 horses we observed a significant effect of the *MC1R-ASIP* signaling system on the APHA-defined Regular registry white spotting phenotype (Table 4, p = 0.0171), with the *e/e* genotype producing the highest proportion of Regular registry horses.

The sex of the individual horse did not significantly impact the registry designation when examined within 461 horses possessing just one white spotting allele (Chi-square test, p = 0.209). Across the entire sample population of 1,054 horses, stallions were more likely to enter the Regular registry than mares or geldings (p< 0.0001), but this may reflect a tendency for applicants to submit registrations on colts with some prospect as future stallions, rather than an influence of sex or castration on white spotting phenotype.

### The KIT*W20 allele results in white spotting, independent of other known alleles

We investigated association between *KIT*W20*, a common allele in the APHA population, and the APHA Regular registry phenotype using a subset of the sample comprised of 529 horses possessing only the *N/N* (n= 270)*, W20/N* (n= 225) and *W20/W20* (n= 34) genotypes (no other spotting alleles at any of the 8 remaining loci tested, to control for potential interactions with other variants.) *KIT*W20* is indeed significantly associated with white spotting patterns on the coat, as defined by the Regular registry APHA threshold (Chi-square test under a dominant model, p= 1.86×10^−18^, Table 3). Clearly, *KIT*W20* strongly correlates with the white spotting phenotype as defined by the *APHA Rule Book.* Novel white spotting patterns likely appear in 2.9% of the population (discussed above), and thus would not be sufficient on their own to coincidentally explain the effect of *KIT*W20* on the Regular registry spotting phenotype, as has been previously suggested by breeders of APHA horses. The significantly increased likelihood of achieving Regular registry status among horses carrying only the *KIT*W20* allele conclusively demonstrates the value of this trait in genomic selection schemes for breeders seeking to improve their production of white spotted, Regular registry eligible foals.

### No evidence for unknown white spotting phenotype “modifying” alleles

Given the historical selective pressure to eliminate white spotting phenotypes in the American Quarter Horse breed (AQHA 2019) and the rarity of white spotting phenotypes within the Thoroughbred breed (often used in the production of American Quarter Horses), APHA breeders are concerned that undiscovered “modifying” alleles capable of reducing the white spotting phenotypes may be present in the American Quarter Horse population. As APHA rules permit cross-breeding with American Quarter Horses or Thoroughbred horses, as well as registration of horses from these breeds that exhibit approved white spotting phenotypes, in-flow of “modifying” alleles could negatively impact the expression of white spotting phenotypes on APHA horses in future generations. 335 registrations were examined where the sire of the submitted horse belonged to the APHA Regular registry and the dam was registered as either Solid Paint-Bred (n= 189) versus those with dams from the American Quarter Horse Association or The Jockey Club (Thoroughbred) registries (n= 146). No significant difference was observed in the Regular vs. Solid Paint-Bred registry status of foals from dams belonging to the Solid Paint-Bred registry compared to those with dams registered with the AQHA or The Jockey Club (Chi-square test p= 0.704). Finally, at the genome-wide scale, the genetic background of horses from the APHA and AQHA registries is very similar (Petersen et al. 2013), and written history documents many AQHA-registered foundation animals at the inception of the APHA.

## Conclusions

Given the large population size of the APHA, estimated at over a million animals, our sampling of just 1,054 animals may not be large enough to accurately reflect allele frequencies across the breed as a whole. However, this is the largest sampling of this breed accomplished to date and provides a first glimpse into the potential value of genotype assisted selection for spotting pattern in the APH. Here we document that most of the genetic markers responsible for the iconic color patterns valued by American Paint Horse Association members, breeders and owners. These loci are valuable tools for prediction of the genetic value of breeding stock, thereby optimizing future production of spotted foals. Identification of phenotype-altering interactions between the nine spotting loci tested, as well as with the *MC1R-ASIP* signaling system, will improve genotype-based phenotype predictions. Genetic tools could rapidly improve the accuracy of selection for white spotting in the horse and may provide significant economic savings compared to the time-consuming photo-analysis approach currently used to register APHA horses.

## Acknowledgments

The authors would like to thank the many APHA staff members for their efforts in submitting and collating the data analyzed in this study. Thanks to the UF undergraduate researchers who generously volunteered for data-entry work on this project: Hannah Hillard, Kalisse Horne, Rachel Kullman, Erica Riano, Matt Winter, Courtney McCreary, Rachel Shepherd, Anna Moskovitz, and Kaycie Miller. Our gratitude to Dr. Ernie Bailey for proofreading the manuscript.

