## Supplemental Figures/Tables for "Impact of white spotting alleles, including *W20*, on phenotype in the American Paint Horse"

**Figure/table S1 (graphs with embedded tables follow):**

Total horse counts and distribution for demographic factors among the sample of 1054 American Paint Horses, colored by registry designation to the Regular (colored with a diagonal pattern) or Solid Paint-Bred sub-registry (light-green colored bars). The four factors presented here are: a) sub-registry, b) sex, c) age when submitted for registration and d) year registration granted.

a) **Sub-registry**

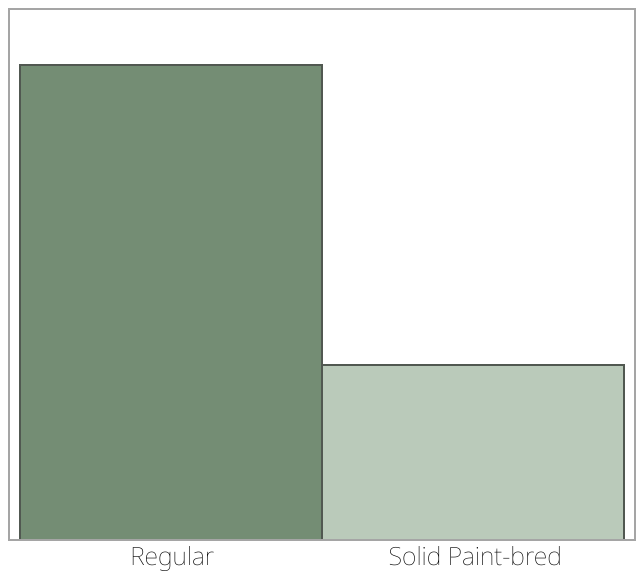

**Frequencies**

| **Level** | **Count** | **Prob** |
| --- | --- | --- |
| Regular | 773 | 0.73340 |
| Solid Paint-bred | 281 | 0.26660 |
| Total | 1054 | 1.00000 |

**b) Sex (F= Female, G= Gelding, M= Male)**

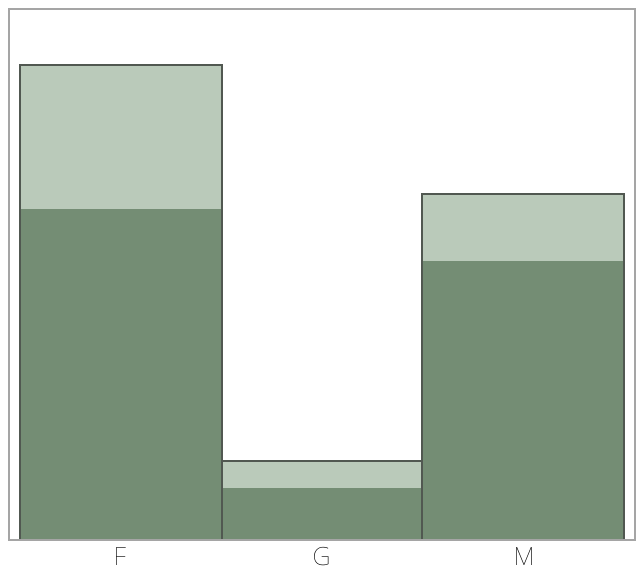

**Frequencies**

| **Level** | **Count** | **Prob** |
| --- | --- | --- |
| F | 558 | 0.52941 |
| G | 90 | 0.08539 |
| M | 406 | 0.38520 |
| Total | 1054 | 1.00000 |

b) **Age at Registration**

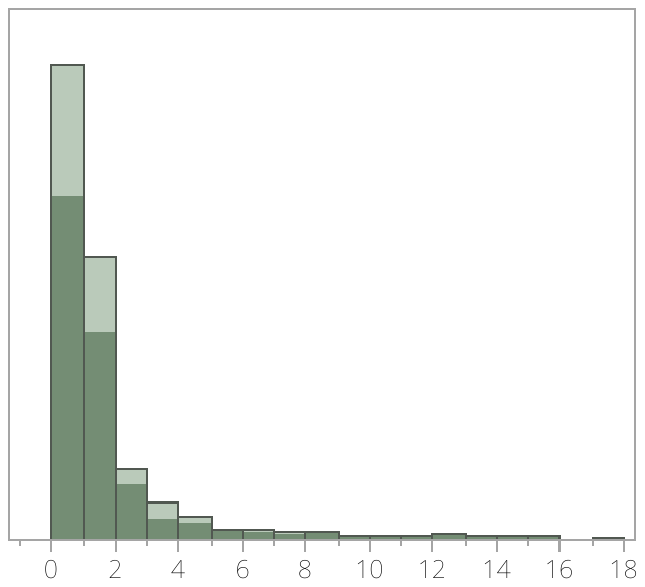

**Summary Statistics**

| Mean | 1.0749526 |
| --- | --- |
| Std Dev | 2.0527421 |
| Std Err Mean | 0.0632287 |
| Upper 95% Mean | 1.1990211 |
| Lower 95% Mean | 0.950884 |
| N | 1054 |

c) **Registration Year**

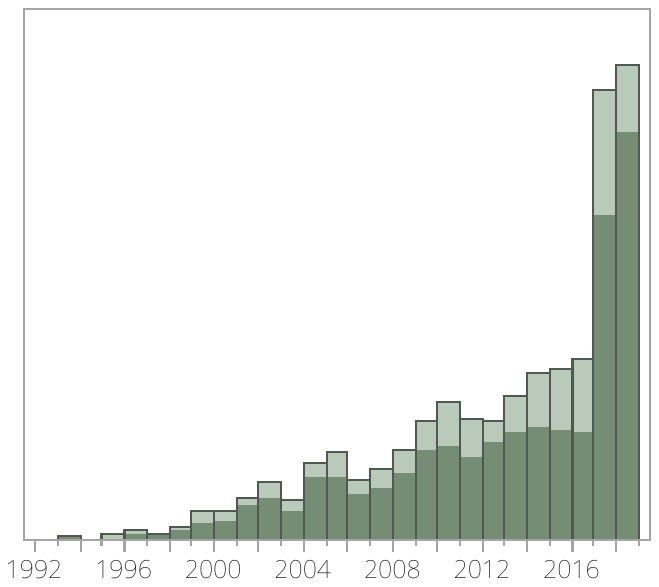

**Summary Statistics**

| Mean | 2012.4858 |
| --- | --- |
| Std Dev | 5.4332946 |
| Std Err Mean | 0.1673566 |
| Upper 95% Mean | 2012.8142 |
| Lower 95% Mean | 2012.1574 |
| N | 1054 |
